# Genome-wide dissection of tillering responsiveness to neighbour proximity in sorghum

**DOI:** 10.64898/2026.07.08.737219

**Authors:** Asad Riaz, Sofie Pearson, Colleen Hunt, Sivakumar Sukumaran, Yongfu Tao, Mark Cooper, Graeme Hammer, Emma Mace, David Jordan

**Affiliations:** Centre for Crop Science, Queensland Alliance for Agriculture and Food Innovation, The University of Queensland, St Lucia, QLD, 4072, Australia; The ARC CoE for Plant Success in Nature and Agriculture, The University of Queensland, St Lucia, QLD, 4072, Australia; Agri-Science Queensland, Department of Agriculture and Fisheries (DAF), Hermitage Research Facility, Warwick, QLD, 4370, Australia; Grains Innovation Park, 110 Natimuk Road, Agriculture Science and Technology, Agriculture Victoria, Horsham, Victoria, 3400, Australia; State Key Laboratory of Tropical Crop Breeding, Shenzhen Branch, Guangdong Laboratory of Lingnan Modern Agriculture, Key Laboratory of Synthetic Biology, Ministry of Agriculture and Rural Affairs, Agricultural Genomics Institute at Shenzhen, Chinese Academy of Agricultural Sciences, Shenzhen, Guangdong 518120, China

**Author notes:** Corresponding author: Asad Riaz.

**Keywords:** sorghum, tillering plasticity, genome-wide association study (GWAS), neighbour detection, red:far-red light, plant density

## Abstract

Tillering plasticity is a key adaptive trait in sorghum influencing resource use efficiency via a plant’s ability to adjust branching to neighbour density. Neighbour detection through red:far-red (R:FR) light sensing regulates this plasticity. While molecular pathways regulating tiller outgrowth are partly known, the genetic architecture underlying density-responsive tillering has not been resolved in any grass species. A sorghum diversity panel (n = 895) was evaluated over two growing seasons (2023 and 2024) with plant spacing ranging from 5 to 60 cm. A linear mixed model incorporating neighbour distance and tiller counts estimated genotype-specific response. GWAS was conducted on isolated plants (no neighbours within 60 cm) and on estimated responsiveness to neighbours. GWAS identified 52 baseline tillering QTLs and 50 for spacing responsiveness, with 10 overlapping, suggesting shared genetic control. Comparison with 41 R:FR pathway candidate genes revealed enrichment in responsiveness QTLs (5/50, 10%) versus baseline (0/52, 0%) (Fisher’s exact test, P = 0.025). Our model identified 40 unique density-responsive tillering QTL regions. Reducing genotype response to neighbour absence could be a selection target to develop water-efficient sorghum varieties where controlled architecture may be more valuable than natural plasticity.

## 1. Introduction

Tillering plasticity, the ability of cereal crops to adjust their branching architecture in response to environmental cues, is a pivotal trait contributing to grain yield and resource use efficiency under varying planting density conditions. In sorghum (*Sorghum bicolor* (L.) Moench), a drought tolerant cereal critical for global food security, tillering (the production of lateral shoots or tillers from the basal node) impacts grain number and canopy size, and shapes crop performance across diverse agroecosystems (Kebrom & Mullet, 2015; Borrell *et al.,* 2000). Tillering plasticity is particularly crucial in water limited environments, where efficient tillering optimises resource allocation, enhancing water use efficiency and sustaining yield under agricultural drought stress (Casal & Fankhauser, 2023; Vadez *et al.,* 2024). One of the primary mechanisms driving tillering plasticity is neighbour detection, mediated by phytochrome photoreceptors that sense changes in the red:far-red (R:FR) light ratio. Under crowded conditions, neighbouring plants absorb red light for photosynthesis while reflecting far-red light, creating a characteristic low R:FR environment that signals competitive pressure and triggers shade avoidance responses, including reduced tillering (Casal *et al.,* 1986; Holmes & Smith, 1975; Ballaré *et al.,* 1987).

Phytochromes, phytochrome B (PHYB) and phytochrome A (PHYA), are the primary sensors of R:FR changes in sorghum, regulating neighbour detection and shade avoidance responses (Feldman *et al.,* 2017; Martínez-García *et al.,* 2010). PHYB is the dominant photoreceptor, transitioning between its inactive (Pr) and active (Pfr) forms based on R:FR ratios. Under low R:FR conditions indicative of high plant density, PHYB accumulates in its inactive Pr form, triggering downstream responses that inhibit axillary bud outgrowth and reduce tillering (Kebrom *et al.,* 2010; Kebrom & Mullet, 2016). In model species such as *Arabidopsis thaliana*, extensive studies have identified key regulators of shade avoidance. The COP1 gene, encoding a ubiquitin E3 ligase, modulates neighbour detection by degrading transcription factors like HY5 under low R:FR conditions (Deng *et al.,* 1991; Casal, 2013). In sorghum, SAV4 has been shown to enhance shade avoidance responses by integrating PHYB signals, suppressing tillering in dense canopies (Sebastian *et al.,* 2020). Conversely, in low planting density with high R:FR ratios, PHYB’s active Pfr form promotes tillering (Kebrom & Brutnell, 2007; Finlayson *et al.,* 2010). PHYA complements PHYB by enhancing shade avoidance under deep canopy shade, where R:FR ratios are extremely low, ensuring robust responses to varying density gradients (Martínez-García *et al.,* 2010). Phytochrome interacting factors (PIFs) integrate these signals, regulating SAV4 and other downstream genes to modulate tillering in response to neighbour proximity (Leivar & Quail, 2011).

The physiological and genetic basis of tillering in sorghum has been progressively dissected through a series of key studies. Foundational work by Lafarge & Hammer (2002) first established the hierarchical dynamics of tiller emergence and fertility across a density gradient, providing a crucial physiological framework. Building on this, Alam *et al.,* (2014a) developed a comprehensive physiological model and identified quantitative trait loci (QTLs) for tillering, primarily under non-competitive conditions Alam *et al.,* (2014b). Their subsequent work improved the prediction of tillering across environments (Alam *et al.,* 2017), and recent modelling efforts continue to refine our understanding of tiller dynamics (Hammer *et al.,* 2023). However, while these studies have been instrumental in characterizing tillering propensity (inherent tiller production under non-competitive conditions; Alam *et al.,* 2014a), they have not systematically mapped the genetic architecture underlying the plastic response to neighbour proximity itself. Despite these molecular insights, the genetic architecture of R:FR mediated tillering plasticity in sorghum, remains poorly characterised compared to model species like Arabidopsis and rice. In model species, extensive studies have identified key regulators of shade avoidance including *PhyB*, *PhyA*, *COP1*, *PIFs*, and *SAV4* which govern light signalling responses (Leivar & Quail, 2011; Kikuchi *et al.,* 2017). In contrast, sorghum research has identified only a few regulators such as *PhyB* and *PIF* genes, leaving a critical gap in understanding the genetic basis of neighbour induced tillering plasticity (Wang *et al.,* 2022; Sebastian *et al.,* 2020). Prior studies have focused on tillering propensity rather than neighbour-induced plasticity, which is essential for field performance under varying plant density (Sawers *et al.,* 2005). To date, no genome-wide association studies have explicitly mapped loci controlling genotypic responsiveness of tillering to plant density in grasses. Instead, most GWAS have focused on tiller number or plant architecture measured at single densities, limiting mechanistic insight into plastic responses relevant for high-density or water-limited environments (Kikuchi *et al.,* 2017). This gap is compounded by the complexity of tillering, influenced by multiple genes and environmental factors, and the challenges of phenotyping density dependent responses in field conditions (Feldman *et al.,* 2017).

This study presents the first GWAS in sorghum to explicitly investigate the genetic basis of neighbour-dependent tillering responses, leveraging extensive genetic diversity to identify loci associated with tillering plasticity. We imposed continuous plant spacing treatments (5-60 cm) across two growing seasons to induce different R:FR gradients in field conditions where neighbour proximity alters light quality and drives shade avoidance responses. Using a linear mixed spatial model, we quantify genotype specific tillering responses to neighbour proximity, which capture linear and linear spatial effects unique to each genotype. While these natural neighbour detection mechanisms evolved to optimise plant survival in variable environments, their agricultural utility may be more complex. In modern farming systems, predictable plant architecture often holds greater importance than natural plasticity, particularly where consistent planting densities and controlled agronomic environments are maintained. Conversely, in variable or low-input systems, maintaining tillering plasticity may allow genotypes to better exploit available resources when neighbour competition is reduced (Alam *et al.,* 2014a). The objectives of this study are to: (1) quantify the relationship between neighbour distance and tillering under varying R:FR ratios induced by plant density, and (2) identify quantitative trait loci (QTL) associated with light sensing genes controlling neighbour detection responses in sorghum. By understanding the genetic basis of natural plasticity responses, breeding strategies can be designed to either harness this plasticity when beneficial or suppress it when agricultural optimization requires more predictable plant architectures. This approach will enhance yield stability and resource efficiency under both controlled agronomic management and variable field conditions, addressing the challenges of climate change and supporting global food security. By identifying genetic regions controlling neighbour-induced tillering, this study establishes a molecular framework to inform the development of sorghum varieties that more effectively enable leveraging of seasonal predictions to produce optimal canopy sizes under high-density and water-limited conditions. Improved understanding of neighbour detection pathways has the potential to enhance yield stability and resource-use efficiency, contributing to climate-resilient cropping systems and long-term food security (Casal & Fankhauser, 2023).

## 2. Materials and methods

### 2.1. Field trials and plant material

Field experiments were established at Hermitage Research Facility, Warwick, Queensland, Australia (28°12’S, 152°5’E, 470 m above sea level) during two consecutive growing seasons (2023 and 2024). A diversity panel of 895 sorghum inbred lines (genotypes), representing global genetic diversity, was used for this study (Tao *et al.,* 2020). Trials were planted using a row-column design with two replicates per year. Each plot consisted of 3 meter long rows with 0.75 m row-to-row spacing. A total of 894 genotypes were evaluated in the first season and 887 in the second season, including 886 genotypes common across both years. The 2023 trial used a 26 × 72 row-column layout, and the 2024 trial used a 36 × 50 layout to accommodate the available genotypes. Standard agronomic practices were followed, including pre-plant fertilizer application and routine pest monitoring and control measures. Irrigation was applied during establishment to ensure uniform germination, with trials managed under rainfed conditions thereafter.

### 2.2. Plant spacing treatments for tillering plasticity

To create varying neighbour density environments, all plots were initially sown at approximately 10 cm spacing. Post-emergence, natural variation in plant establishment resulted in within-row spacings ranging from approximately 5 to 20 cm. Focal plants were then selected and classified into three density categories: isolated, medium density and high density. For isolated treatments, surrounding plants within 60 cm were removed by hand-thinning before the three-leaf stage to eliminate neighbour interactions. This distance threshold was informed by previous field observations indicating that neighbour effects on tillering become negligible beyond approximately 60 cm spacing (Lafarge & Hammer, 2002). For the density treatments, medium density focal plants were spaced 20-40cm from neighbouring plants, while high density focal plants were spaced 5-20cm from neighbours. This approach created three distinct competitive environments while maintaining realistic field conditions for the density treatments and providing continuous datasets. Given that the row-to-row spacing of 0.75 m exceeded the 60 cm threshold at which neighbour effects on tillering become negligible, between-row neighbour interactions were not considered and neighbour distance measurements focused on within-row spacing. Individual plant positions and neighbour distances were measured with a ruler to quantify the local competitive environment for each focal plant. Distance measurements to neighbours on both sides were recorded, with the shorter distance classified as the nearest neighbour and the longer distance as the farthest neighbour to capture the full spatial context affecting each plant.

### 2.3. Phenotyping and data collection

Neighbour distance measurements and tiller counts were recorded simultaneously at physiological maturity, so focal plants did not require physical marking. Isolated plants were located at one end of each plot, identifiable by the absence of neighbours within the hand thinned zone. Tiller number was recorded at physiological maturity for each focal plant by counting all tillers visible on the plant, including both green and senesced tillers. Shoots emerging from axillary buds at the basal nodes that had developed beyond the three-leaf stage were counted as tillers. This count may not include tillers that initiated early in development, senesced, and were therefore no longer visible at maturity (Lafarge & Hammer, 2002). For isolated plants, Best Linear Unbiased Estimators (BLUEs) were calculated to extract adjusted phenotypic values accounting for spatial field variation. BLUEs were computed using ASReml-R v.4.2 (Butler *et al.,* 2009) in R v.4.4.1 (R Core Team, 2024) with a linear mixed model incorporating row and column positions as random effects with autoregressive spatial correlation structure to control for field heterogeneity (Gilmour *et al.,* 1997).

### 2.4. Genotype filtering strategies for responsiveness analysis

A subset of genotypes produced no tillers or very few tillers at low densities (ie the isolated treatment) making them uninformative for assessing the response to density even though they potentially contained genes for density responsiveness. The inclusion of the non-informative genotypes would reduce the power of QTL analyses. To address this issue, genotypes in the isolated treatment datasets for each site were removed using multiple filtering strategies depending on their tillering capacity. Genotypes were progressively filtered based on their tiller number in the isolated treatment prior to spatial modelling. We evaluated four filtering strategies using the isolated plant treatment: (1) Basic; ≥1 Tiller (retain all genotypes that produced at least one tiller), (2) >1 Tiller, (3) >2 Tillers, and (4) >3 Tillers. The corresponding sample sizes for each filtered dataset are detailed in Table 1. The impact of these filters on QTL detection was assessed subsequently. The four filtering strategies were evaluated by inspecting QQ plots and genomic inflation factors (λ) of the resulting GWAS analyses to assess test statistic inflation. λ values were comparable across all filters and consistent with well-controlled population structure, indicating that filtering did not introduce systematic bias. The final filter was therefore selected based on QTL detection power rather than inflation control.

**Table 1.** Number of genotypes retained after progressive filtering.

| Year | Filter | No of Genotypes |
| --- | --- | --- |
| 2023 | Basic ($\geq 1$ Tiller) | 830 |
| 2024 | Basic ($\geq 1$ Tiller) | 827 |
| Combined | Basic ($\geq 1$ Tiller) | 775 |
| 2023 | >1 Tiller | 729 |
| 2024 | >1 Tiller | 545 |
| Combined | >1 Tiller | 493 |
| 2023 | >2 Tiller | 633 |
| 2024 | >2 Tiller | 484 |
| Combined | >2 Tiller | 325 |
| 2023 | >3 Tiller | 420 |
| 2024 | >3 Tiller | 353 |
| Combined | >3 Tiller | 153 |

Combined: genotypes common to both growing seasons (2023 and 2024) that met the filtering criterion.

### 2.5. Spatial modelling and responsiveness quantification

Following genotype filtering (section 2.4), responsiveness to neighbour spacing was quantified for the retained genotypes in each filtered dataset using a linear mixed model that captured individual tillering responses to plant spacing:

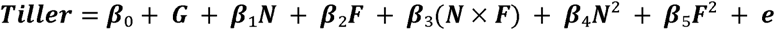

Where:

- âL = overall site mean (baseline tillering level)
- *G* = genotype-specific main effect (inherent tillering capacity differences)
- *N* = nearest neighbour distance
- *F* = farthest neighbour distance
- âL = genotype-specific response to nearest neighbour distance
- âL = genotype-specific response to farthest neighbour distance
- âL = interaction between nearest and farthest neighbour distances
- âL, βL = quadratic terms modelling non-linear spacing responses
- *e* = residual error

This model was applied to each genotype to derive individual spatial response coefficients (β_-β_), quantifying genotype-specific responsiveness to neighbour proximity. The quadratic terms captured non-linear responses such as diminishing light effects as spacing increased. To standardize responsiveness quantification across genotypes, the derived spatial coefficients were used to predict tillering responses at equidistant neighbour configurations. Predicted tiller numbers were calculated at standardized distances of 10-10 cm, 20-20 cm, and 30-30 cm (nearest-farthest neighbour distances). Responsiveness was quantified as the slope difference between 10-10 cm and 20-20 cm predictions, representing the rate of tillering change per unit distance increase. This standardized responsiveness metric was used as the quantitative trait for subsequent analysis.

### 2.6. Genotyping and SNP data

Genotyping was performed as described in Tao *et al.,* (2020) using the same diversity panel. Briefly, DNA was extracted from bulked young leaves using the CTAB method and genotyped using DArTseq technology. Following quality control filtering for minor allele frequency >0.01 and missing data <50%, the final dataset comprised 742,000 SNPs across the diversity panel for GWAS analysis.

### 2.7. Genome-wide association analysis

GWAS was performed using FarmCPU (Liu *et al.,* 2016). The genome-wide significance threshold was determined using the simple M method (Li & Ji, 2005), which estimates the effective number of independent tests by accounting for linkage disequilibrium between SNPs. Based on this correction, a threshold of *P* < 1×10□□ (−log□□ *P* > 5) was applied across all GWAS analyses to declare SNPs genome-wide significant. Separate GWAS analyses were conducted for baseline tillering (tiller number of isolated plants without neighbour effects, representing inherent tillering capacity) and neighbour responsiveness traits. Linkage disequilibrium analysis revealed that LD decayed to background levels at approximately 1 cM in the diversity panel . Therefore, significant SNPs were clustered into quantitative trait loci (QTL) using a 2-cM window approach (1 on each side), as described in Tao *et al.,* (2020). SNPs within 2 cM of each other were grouped into single QTL regions based on genetic map positions from the sorghum consensus map (Mace *et al.,* 2009, 2019). Within each QTL region, the SNP with the lowest *P*-value was designated as the peak SNP and used to represent the QTL position in summary tables. Where multiple independent significant SNPs occurred within the same 2-cM window, all SNPs were retained and reported in the supplementary SNP table, while the QTL was represented in the main results by its peak SNP.

### 2.8. QTL overlap and co-localization analysis

Overlap between baseline tillering QTL (isolated plants) and neighbour responsiveness QTL was assessed using a 2-cM window approach. QTL were considered overlapping if their peak SNPs were within ±1 cM of each other. For each QTL identified in any analysis, the −log□□(*P*) of the peak SNP was extracted from all six GWAS analyses (BL_23, BL_24, BL_com, R_23, R_24, R_com) and reported in Table 4 to enable cross-analysis comparison of signal strength. Only values exceeding the significance threshold were considered detected QTLs in that specific analysis; sub-threshold values are reported for transparency and to assess concordance of signal across analyses.

Co-localization analysis was performed to test whether identified QTL regions were associated with R:FR light signalling and neighbour detection pathways. A total of 41 sorghum orthologues of candidate genes for R:FR light perception, phytochrome signalling, shade avoidance, and R:FR-regulated branching responses were compiled from Arabidopsis and rice and mapped to their genetic positions (Table S1). Candidate genes were considered co-localized with a QTL if the gene’s position on the sorghum consensus genetic map (Mace *et al.,* 2009, 2019) was within ±1 cM of the peak QTL SNP. To test whether neighbour responsiveness QTLs were enriched for R:FR pathway candidate genes relative to baseline tillering QTLs, Fisher’s exact test (alternative = “greater”) was conducted on a 2×2 contingency table with rows representing the two QTL sets (responsiveness QTLs, *n* = 50; baseline QTLs, *n* = 52) and columns representing colocalization status (colocalized with a candidate gene, not colocalized). The test evaluated the null hypothesis that the proportion of QTLs colocalized with R:FR candidate genes did not differ between the two sets. Analyses were conducted in R v.4.4.1 (R Core Team, 2024) using the fisher.test() function.

## 3. Results

### 3.1. Inducing neighbour detection responses through plant spacing treatments

Plant spacing treatments were categorised as three density categories across the sorghum diversity panel: isolated (farthest neighbour >60 cm), medium density (nearest neighbour 20-40 cm), and high density (nearest neighbour 5-20 cm). In the 2023 trial, these treatments induced strong tillering responses, with high-density conditions showing significant tillering reduction compared to isolated plants (mean 1.5 vs 3.7 tillers per plant, 60% reduction; *P* < 0.0001; Fig.S1). Medium density treatments produced intermediate responses with a mean of 2.3 tillers per plant in 2023, representing a 38% reduction compared to isolated conditions (*P* < 0.0001; Fig. S1). Similar patterns were observed in 2024, with high-density conditions showing significant tillering reduction compared to isolated plants (mean 1.5 vs 3.3 tillers per plant, 53% reduction; *P* < 0.0001) and medium density treatments producing intermediate responses (mean 2.4 tillers per plant, 27% reduction; *P* < 0.0001; Fig. S1). Summary statistics for both years are presented in Table S2.

Fig. 1 (a-b) suggest that there is substantial genetic variation in tillering response to density among the genotypes with some genotypes showing similar tiller numbers in high and medium density situations compared to the isolated plants while others showed substantial reduction in tillering in response to neighbours. These qualitative results were supported by the statistical analysis (see below).

**Fig. 1.**
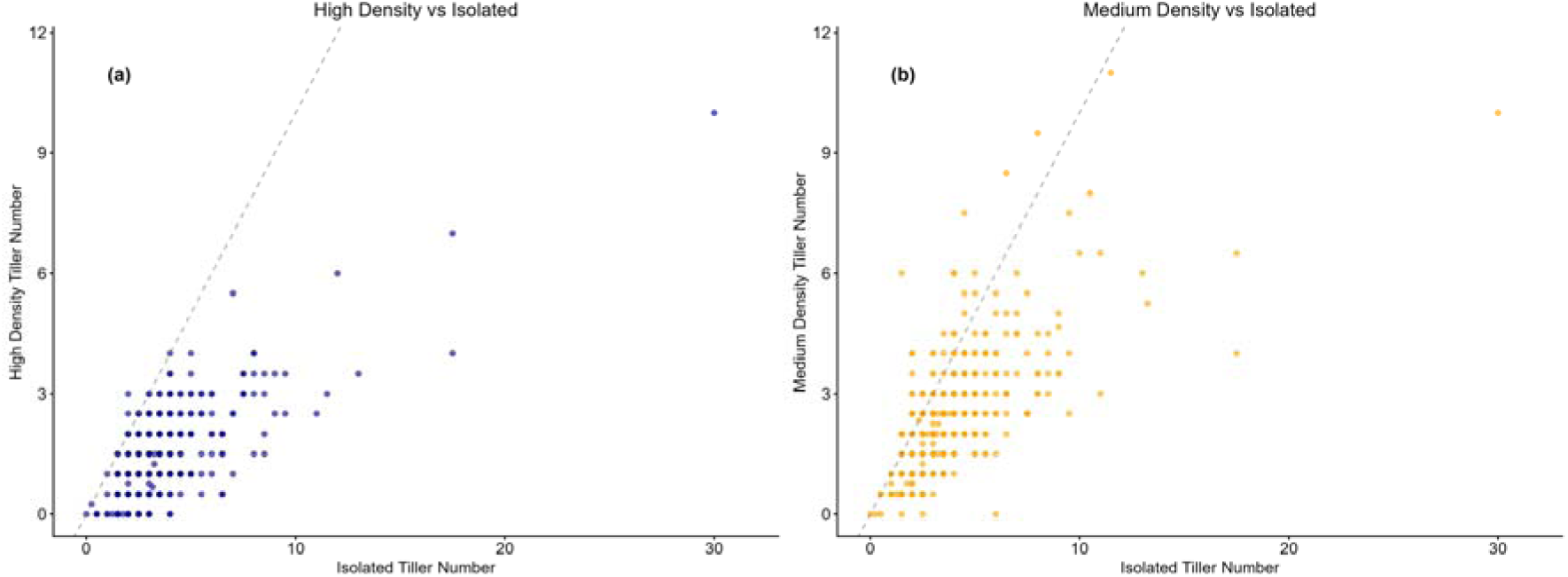
Genotypic variation in tillering responses to density treatments. **(a)** Scatter plot showing the relationship between isolated and high density tillering across genotypes in 2023. **(b)** Scatter plot showing the relationship between isolated and medium density tillering across genotypes in 2023. Each point represents one genotype’s mean tillering response under the respective density conditions. The grey dashed line represents the theoretical 1:1 relationship if genotypes showed no density response. Scatter around this line indicates genetic variation in neighbour sensitivity.

### 3.2. Model performance across sorghum growing seasons

The linear mixed model demonstrated successful fit for quantifying genetic variation in tillering responsiveness to neighbour distance effects across both growing seasons (Table 2). Model convergence was achieved after 12 iterations for both datasets, with final log-likelihoods of −3970.789 (2023) and −4087.466 (2024). Both years showed highly significant genotype main effects (2023: z-ratio = 6.57, *P* < 0.001; 2024: z-ratio = 8.00, *P* < 0.001), confirming substantial genetic diversity in tillering across the sorghum diversity panel. Genetic variation in neighbour responsiveness was significant in both years (*P* < 0.01 in 2023; *P* < 0.05 in 2024), though with contrasting patterns: 2023 exhibited significant nearest neighbour effects (z-ratio = 3.00, *P* < 0.01) and nearest: farthest interactions (z-ratio = 4.21, *P* < 0.001), while 2024 showed significant nearest neighbour effects (z-ratio = 2.08, *P* < 0.05) and farthest neighbour effects (z-ratio = 3.46, *P* < 0.001). Some quadratic terms were constrained to boundary values, indicating that linear and interaction components adequately captured neighbour response patterns. Spatial effects (Column, Row, Replicate) were minimal or non-significant. The significant genotype-by-distance interaction terms in both years provided the necessary genetic variation for subsequent genome-wide association mapping of tillering plasticity to neighbour density effects.

**Table 2.**
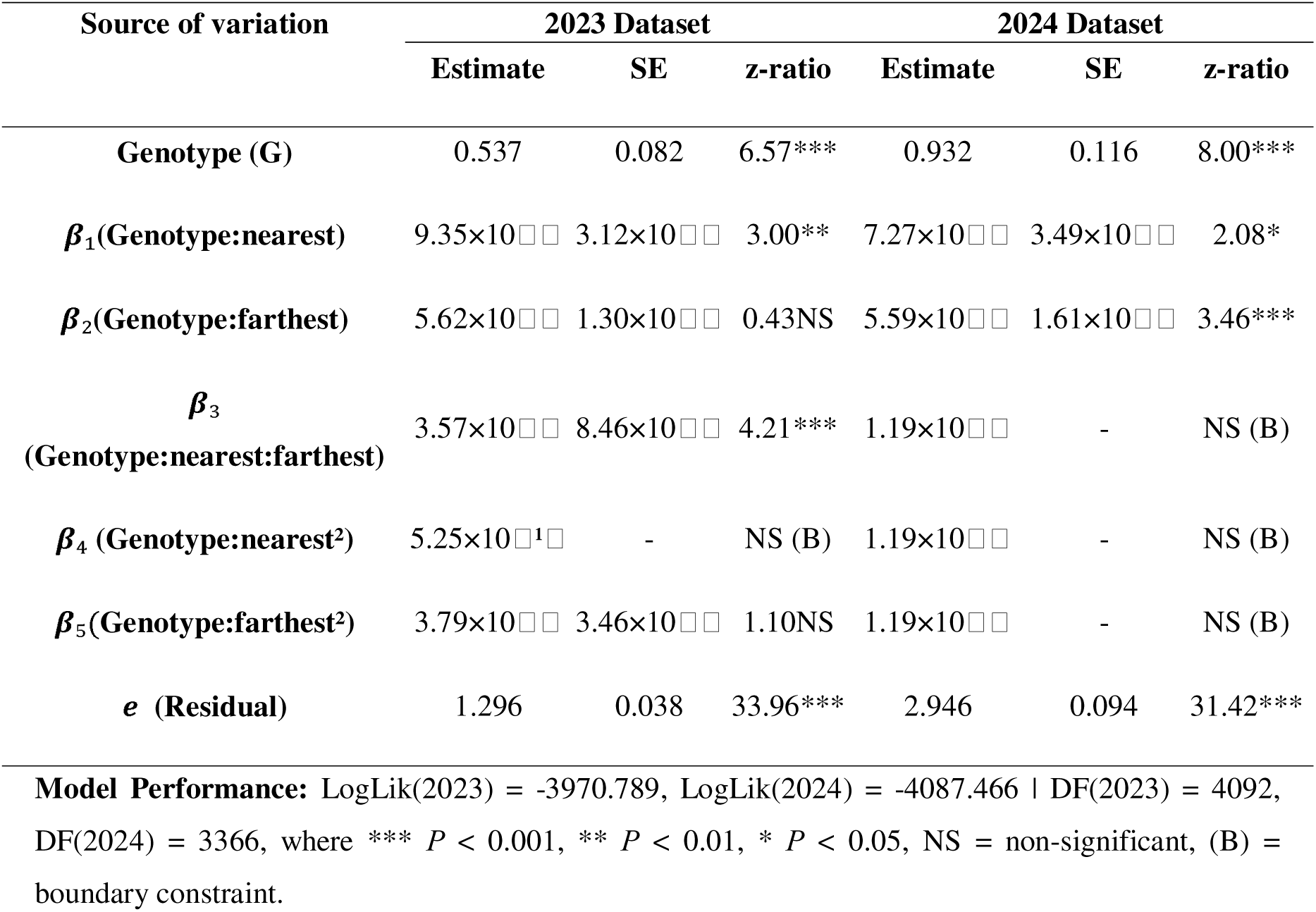
Analysis of variance and statistical parameters for tillering responsiveness model.

### 3.3. Phenotypic variation in baseline tillering and neighbour responsiveness

Substantial variation in baseline tillering capacity (propensity to tiller) was observed across the diversity panel in isolated plants. Baseline tillering exhibited a right-skewed distribution across the diversity panel in 2023, with the majority of genotypes showing low to moderate tillering capacity and a few genotypes exhibiting exceptionally high tillering potential (Fig. 2a). This right skewness was driven by consistent high-tillering genotypes, notably including SC755 that demonstrated reproducible high tiller production across replicates, rather than being an artifact of experimental variation. Pearson correlation of mean tiller number per genotype in isolated plants between 2023 and 2024 was r = 0.629.

**Fig. 2.**
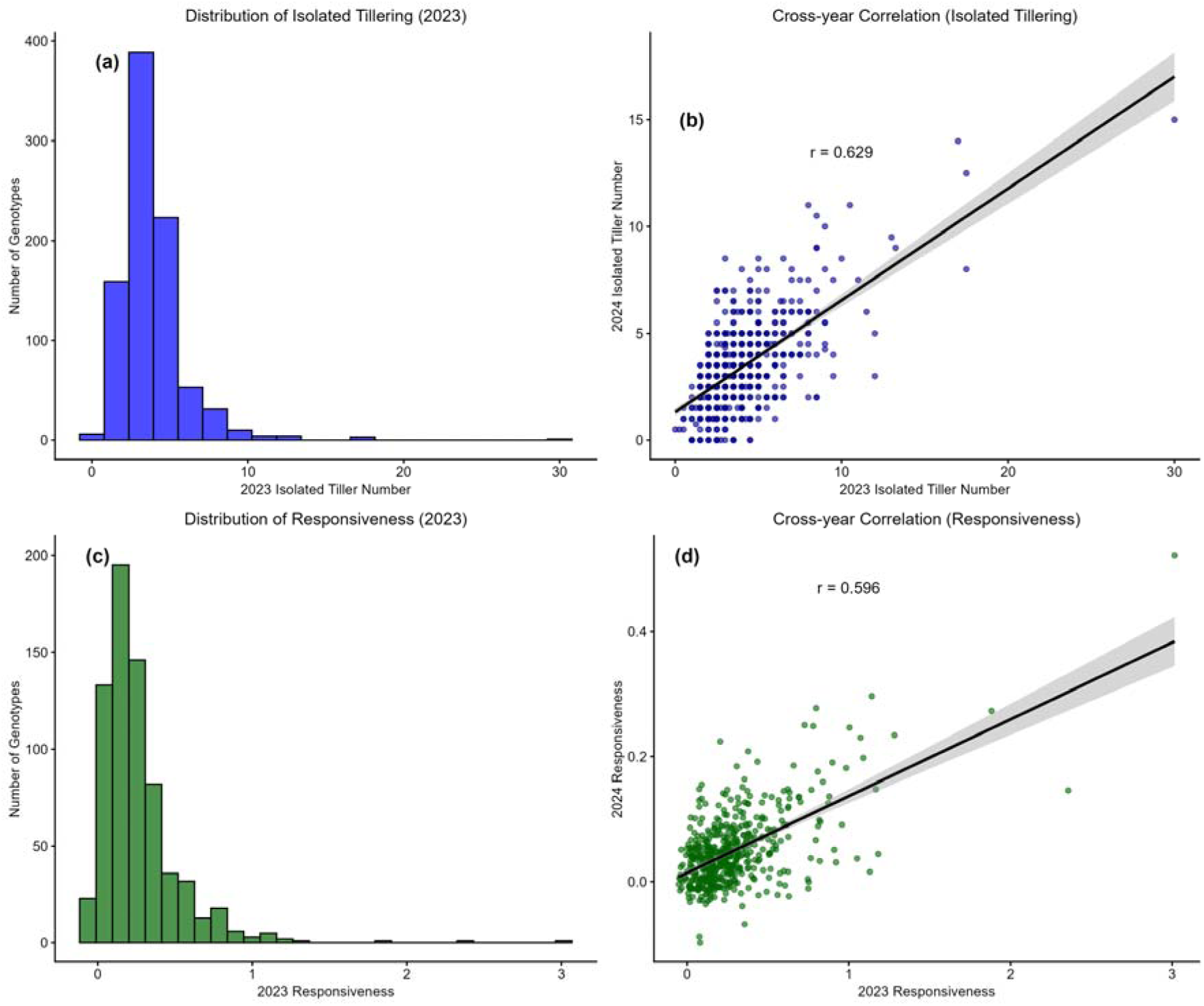
Distribution and stability of tillering traits across years. (a) Frequency distribution of baseline tiller number across genotypes in 2023. (b) Correlation between baseline tiller numbers measured in 2023 (x-axis) and 2024 (y-axis) with Pearson’s r = 0.629. (c) Frequency distribution of tillering responsiveness across genotypes in 2023. (d) Correlation between tillering responsiveness values measured in 2023 (x-axis) and 2024 (y-axis) with Pearson’s r = 0.596. Gray bands in panels (b) and (d) represent confidence intervals (95%) around the linear regression fit (black lines).

Genotype-specific responsiveness to neighbour proximity was successfully quantified using the spatial modelling approach. Neighbour responsiveness in the 2023 trial exhibited a right-skewed distribution ranging from 0 to approximately 3.0 units, representing the standardized slope differences between predicted tillering at 10-10 cm and 20-20 cm neighbour configurations and revealing substantial genetic variation in neighbour detection capacity across the diversity panel (Fig. 2c).

Pearson correlation of genotype responsiveness values (slope difference between predicted tillering at 10-10 cm and 20-20 cm) between 2023 and 2024 was r = 0.596 (Fig. 2d), indicating moderately stable genetic control of neighbour detection mechanisms. The moderate correlation (r = 0.596) compared to baseline isolated tillering (r = 0.631) may reflect differences in the tillering propensity of the environment between years. Additionally, the greater environmental sensitivity of neighbour detection responses, which depend on light quality changes (R:FR) created by neighbour shading that vary with weather conditions and canopy development.

### 3.4. Genetic architecture of tillering plasticity in sorghum

Two distinct GWAS analyses revealed both shared and unique genetic architectures underlying baseline tillering capacity versus neighbour responsiveness in sorghum. GWAS of baseline tillering using BLUEs from isolated plants identified multiple genomic regions contributing to genetic variation across individual years and combined datasets. Single-year analyses detected 21 significant

SNPs in 2023 and 16 SNPs in 2024, while the combined-year analysis yielded 25 SNPs at the genome-wide significance threshold (P < 1×10LJLJ) (Fig. 3). The variation in SNP detection between years reflects sampling error and environmental sensitivity typical of quantitative traits, while the combined analysis provided enhanced power for detecting additional loci not significant in individual years.

**Fig. 3.**
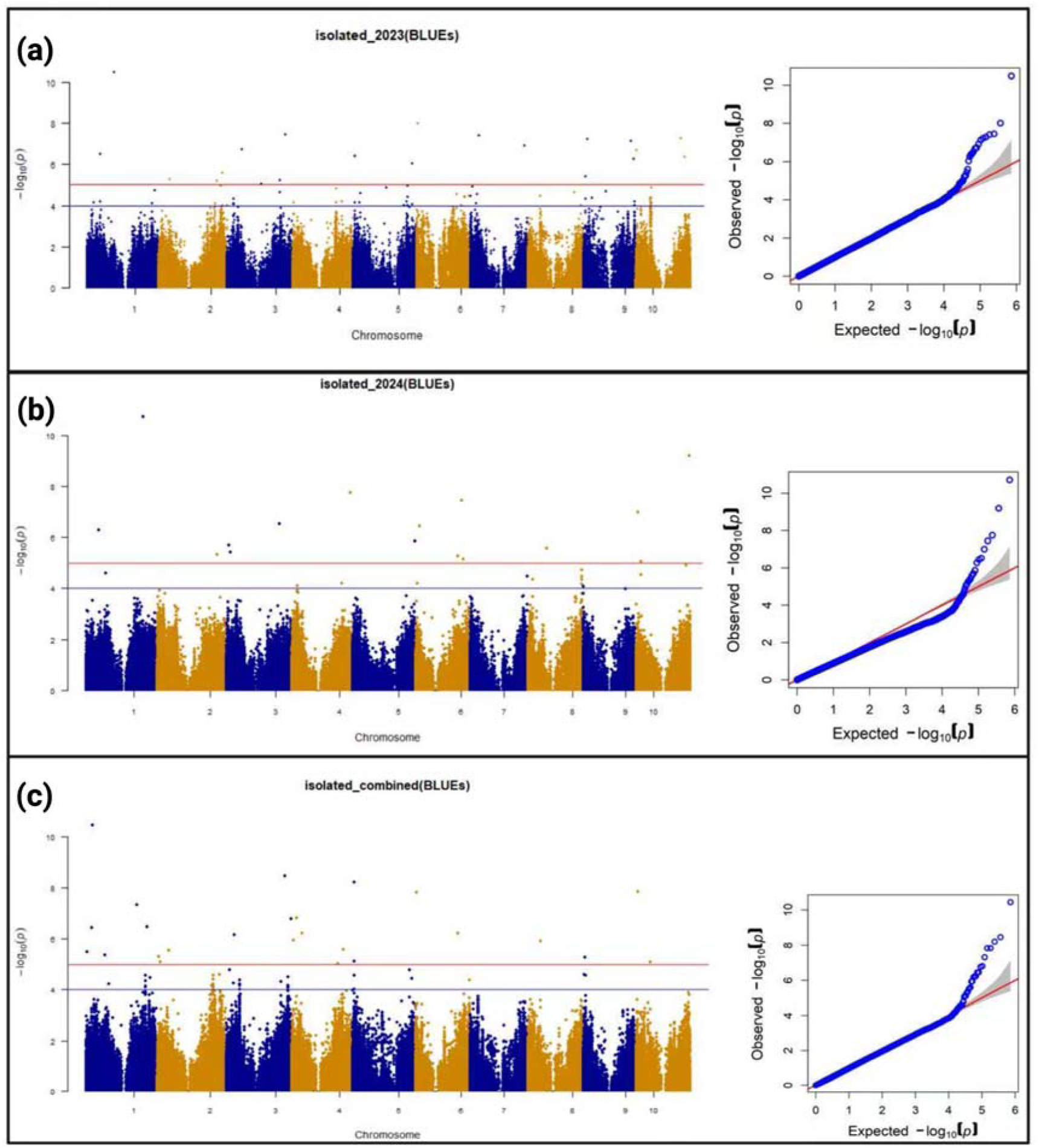
Manhattan and quantile-quantile (Q-Q) plots for baseline tillering GWAS analysis. Manhattan plots showing genome-wide association results for baseline tillering capacity in sorghum across (A) 2023, (B) 2024, and (C) combined years analysis. Physical SNP positions are displayed on the x-axis with alternating blue and orange colours distinguishing chromosomes. The y-axis represents −log□□(P) value of association significance. The horizontal red line indicates the genome-wide significance threshold (P < 1×10□□). SNPs above the threshold represent significant marker- trait associations. Corresponding Q-Q plots show observed (y-axis) versus expected (x-axis) − log□□(P) values, with the red diagonal line representing expected distribution under the null hypothesis and blue points representing observed data. The grey shaded area indicates the 95% confidence interval. Upward deviation from the expected line indicates genuine associations.

Significant marker-trait associations were clustered into QTL regions based on linkage disequilibrium patterns and genetic positions from the sorghum consensus map. A 2-cM window was used to cluster these 62 significant SNPs into 52 unique baseline tillering QTL distributed across all 10 sorghum chromosomes (Table 4), following the approach detailed in Tao *et al.,* (2020). While only SNPs exceeding the genome-wide significance threshold (*P* < 1×10□□) were called as detected QTLs, the *- log*LL*(P)* values of peak SNPs across all six analyses are reported in Table 4, with cell colour intensity reflecting signal magnitude. Several QTLs significant in one analysis showed elevated but sub-threshold signals in additional analyses, indicating concordance of association across years and supporting the consistency of the underlying signals. The 52 baseline QTL were distributed across all 10 sorghum chromosomes, with the highest density on chromosome 1 (9 QTL), with the fewest on chromosomes 7 (2 QTL) and 8 (1 QTL). Among the 52 QTL, 13 regions contained multiple SNPs from different analyses, indicating overlapping signals within these loci, while 39 QTL represented distinct single associations. In the broad genomic distribution, the most significant baseline QTL was qtl1.13 located on chromosome 1 at 64.79 Mb, identified in the 2024 analysis with *P* = 1.89 x 10^-11^. The identification of 52 QTL demonstrates that constitutive tillering capacity in sorghum provides the foundation for understanding how baseline tillering capacity differs from environmentally responsive tillering mechanisms.

GWAS of neighbour responsiveness revealed a more focused genetic architecture targeting specific response mechanisms. Baseline tillering analysis identified the full complexity of tillering biology including resource allocation (sugars), developmental control (phytohormones) and environmental factors, whereas responsiveness analysis captures genes specifically associated with plastic responses to R:FR based neighbour density. This focused phenotype provides enhanced power for QTL detection for this component of tillering despite analysing fewer individuals through filtering approaches. Preliminary analysis of genotypic responsiveness patterns identified four distinct response types: high responsiveness (genotypes showing strong tillering plasticity to neighbour density), moderate responsiveness (intermediate plastic responses), non-informative genotypes (consistently low or zero tiller production), and non-responsive genotypes (constant tiller numbers regardless of neighbour distance) (Fig. 4). The non-informative and non-responsive genotypes were excluded, and the remaining dataset was then subjected to systematic filtering which removed genotypes with varying levels of tillering propensity for reliable neighbour effect measurement (refer to section 2.4). To validate our filtering approach, we compared all strategies in the filtered dataset and selected the No<1 filter (retaining genotypes with >1 tiller) for final GWAS analysis. This decision was based on its performance across three criteria: (1) retention of an adequate sample size while excluding minimally responsive genotypes, (2) optimal QQ plot alignment, and (3) higher unique QTL yield. Together, these results support No<1 as the most biologically relevant choice for analysing tillering responsiveness (Table 3).

**Fig. 4.**
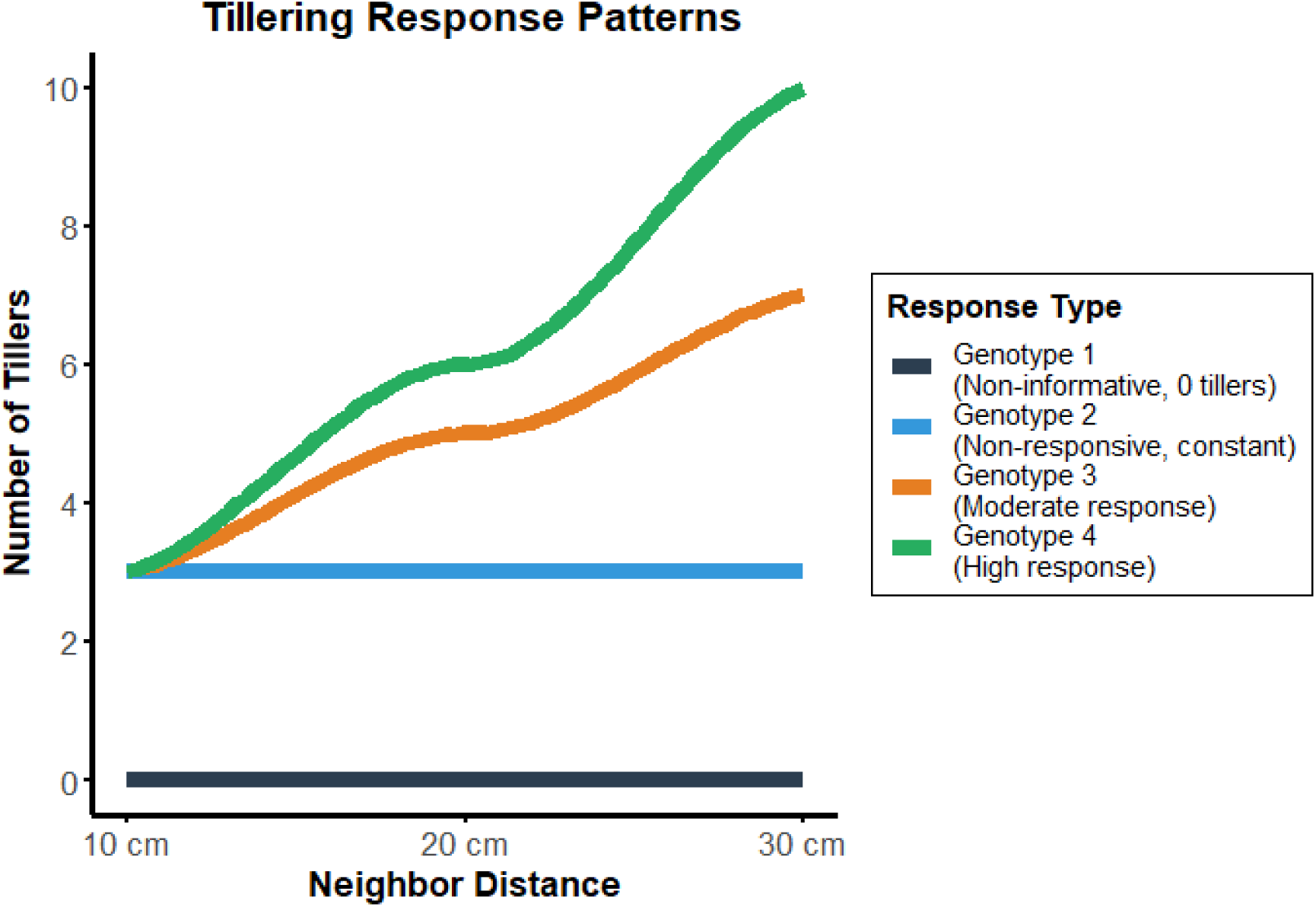
Conceptual framework of tillering response patterns to neighbour distance. Four genotype response types were identified based on tillering plasticity: non-informative genotypes with low tillering (black), non-responsive genotypes with constant (blue) tiller production, and responsive genotypes showing moderate (orange) or high (green) plasticity to neighbour distance changes. Non-informative genotypes were excluded from subsequent GWAS analysis.

**Table 3.** Comprehensive QTL filter comparison.

| Metric | Baseline | No<1 | No<2 | No<3 |
| --- | --- | --- | --- | --- |
| Total Unique QTLs | 52 | 50 | 42 | 39 |
| Baseline Overlaps | - | 10 (20%) | 9 (21.4%) | 8 (20.5%) |
| Unique Plasticity QTLs | - | 40 | 31 | 31 |
| Model Fit (QQ plot) | - | $\lambda \approx 1.0$ | $\lambda < 1.0$ | $\lambda < 1.0$ |

The No<1 filtering approach identified a more comprehensive set of tillering-responsiveness QTLs with the significant peak P-value of 7.05 × 10&#x25Al;¹&#x25Al; (Fig. 5). This dataset identified 65 SNPs, using the same 2-cM clustering approach, these SNPs clustered into 50 unique responsiveness QTLs. Among these QTLs, 40 were unique while 10 were overlapped with baseline tillering showing a minimal overlap (10/50, 20%). Chromosome distribution of responsiveness QTLs showed a different pattern compared to baseline tillering, with chromosome 1 (7 QTLs) followed by 3, 4, 8, and 10 (6 QTLs each), chromosome 6 (5 QTLs), chromosome 5 and 9 (4 QTLs each), chromosome 2 and 7 (3 QTLs each) (Table 4).

**Fig. 5.**
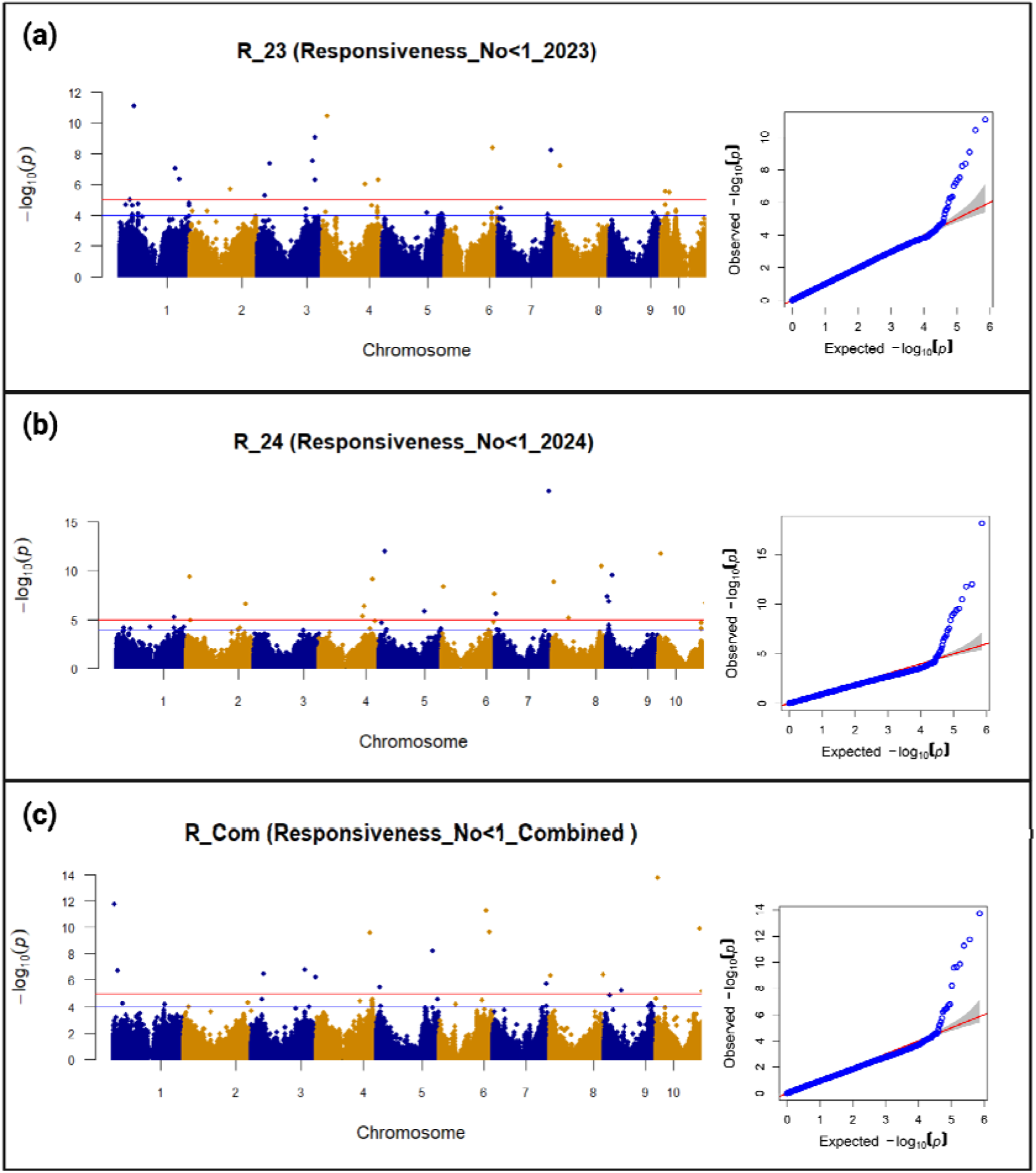
Manhattan and quantile-quantile (Q-Q) plots for neighbour-responsive tillering GWAS analysis using No<1 filtered dataset. Manhattan plots showing genome-wide association results for neighbour responsiveness in sorghum across (A) 2023, (B) 2024, and (C) combined years analysis using the No<1 filtering approach. Physical SNP positions are displayed on the x-axis with alternating blue and orange colours distinguishing chromosomes. The y-axis represents −log□□(P) value of association significance. The horizontal red line indicates the genome-wide significance threshold (P < 1×10□□). SNPs above the threshold represent significant associations with neighbour responsiveness. Corresponding Q-Q plots show observed (y-axis) versus expected (x-axis) −log□□(P) values, with the red diagonal line representing expected distribution under the null hypothesis and blue points representing observed data. The grey shaded area indicates the 95% confidence interval. Upward deviation from the expected line indicates genuine associations, demonstrating enhanced detection power for neighbour-responsive mechanisms compared to baseline analysis.

**Table 4.** Summary of baseline and neighbour-responsive tillering QTLs identified by GWAS in a sorghum diversity panel (*n* = 895) across two growing seasons (2023 and 2024)

| QTLs | Chr | Position (Mb) | Peak (cM) | BL_23 | BL_24 | BL_com | R_23 | R_24 | R_com |
| --- | --- | --- | --- | --- | --- | --- | --- | --- | --- |
| qtl1.1 | 1 | 1.21 | 3.247 | 0.2 | 0.1 | 5.5 | 0.6 | 1.3 | 0 |
| qtl1.2 | 1 | 1.73 | 5.529 | 0.2 | 0.8 | 0.2 | 0.3 | 0.5 | 11.8 |
| qtl1.3 | 1 | 5.5 | 12.853 | 0.1 | 0.2 | 0.2 | 0.9 | 1.3 | 6.7 |
| qtl1.4 | 1 | 6.7 | 16.043 | 3.6 | 1.1 | 6.4 | 2.4 | 0.4 | 1.2 |
| qtl1.5 | 1 | 7.04 | 17.429 | 4.2 | 0.7 | 10.4 | 1.2 | 1.3 | 0.6 |
| qtl1.6 | 1 | 11.37 | 27.779 | 1.3 | 0.6 | 1.5 | 5 | 1 | 0.3 |
| qtl1.7 | 1 | 14.4 | 36.48 | 6.5 | 6.3 | 1.9 | 1.5 | 1.3 | 0.7 |
| qtl1.8 | 1 | 16.98 | 41.41 | 1.2 | 0.2 | 2.3 | 11.1 | 0.8 | 0.7 |
| qtl1.9 | 1 | 21.18 | 46.612 | 0.3 | 0.7 | 5.4 | 0.1 | 0 | 0 |
| qtl1.10 | 1 | 30.45 | 55.315 | 10.5 | 0.6 | 0.1 | 0.5 | 0 | 0.1 |
| qtl1.11 | 1 | 57.22 | 99.1 | 1.4 | 3.2 | 7.3 | 3.7 | 0.1 | 0.5 |
| qtl1.12 | 1 | 64.06 | 132.58 | 1.9 | 0.5 | 1.6 | 7 | 0.6 | 0.2 |
| qtl1.13 | 1 | 64.79 | 136.528 | 0.1 | 10.7 | 0.6 | 0.2 | 0.6 | 2.3 |
| qtl1.14 | 1 | 66.85 | 141.537 | 0.5 | 0 | 0.2 | 0.3 | 5.2 | 1.6 |
| qtl1.15 | 1 | 68.49 | 156.29 | 2.5 | 0.7 | 1.1 | 6.4 | 1 | 0.9 |
| qtl1.16 | 1 | 69.42 | 159.442 | 1.8 | 1.2 | 6.5 | 0.2 | 2 | 0 |
| qtl2.1 | 2 | 1.36 | 1.679 | 0.7 | 1.2 | 5.3 | 1.5 | 0.5 | 0.4 |
| qtl2.2 | 2 | 3.96 | 14.089* | 1.1 | 1.5 | 5.1 | 4.3 | 9.4 | 3.5 |
| qtl2.3 | 2 | 12.76 | 63.226* | 5.3 | 2.1 | 5.6 | 2.9 | 1.1 | 0.6 |
| qtl2.4 | 2 | 46.67 | 75.528 | 0.3 | 0.1 | 1.1 | 5.7 | 1.2 | 0.6 |
| qtl2.5 | 2 | 66.25 | 148.603 | 5.2 | 1.2 | 0.8 | 0 | 1.7 | 0.7 |
| qtl2.6 | 2 | 68.13 | 152.765* | 0.3 | 5.3 | 0.5 | 0.7 | 6.6 | 1 |
| qtl2.7 | 2 | 72.22 | 178.147 | 5.6 | 0.2 | 3.5 | 0.9 | 0.5 | 0.4 |
| qtl3.1 | 3 | 3.33 | 9.627 | 0 | 5.7 | 0 | 0.2 | 1.5 | 0.2 |
| qtl3.2 | 3 | 4.59 | 19.275 | 0.4 | 5.4 | 0.8 | 0.4 | 0.4 | 0.4 |
| qtl3.3 | 3 | 8.90 | 35.052* | 3.6 | 0.1 | 6.2 | 5.3 | 1.9 | 0.6 |
| qtl3.4 | 3 | 14.13 | 50.008 | 1.4 | 1.7 | 3.7 | 7.4 | 2 | 6.5 |
| qtl3.5 | 3 | 17.01 | 56.285 | 6.7 | 1.4 | 2.4 | 1.3 | 0.4 | 0.2 |
| qtl3.6 | 3 | 60.77 | 111.92* | 5.3 | 6.5 | 0.4 | 1 | 1.2 | 1.2 |
| qtl3.7 | 3 | 60.94 | 114.064 | 0.7 | 0.7 | 0.3 | 0.3 | 0.8 | 6.8 |
| qtl3.8 | 3 | 62.61 | 122.456 | 0.5 | 0 | 0.4 | 7.5 | 0.8 | 0.2 |
| qtl3.9 | 3 | 66.19 | 131.51* | 7.4 | 0.3 | 8.5 | 9.1 | 0.5 | 1.3 |
| qtl3.10 | 3 | 73.18 | 163.464 | 0.1 | 1.4 | 0.9 | 1.1 | 1.2 | 6.2 |
| qtl3.11 | 3 | 73.7 | 168.196 | 0.7 | 1.2 | 6.8 | 1.3 | 0.8 | 0.1 |
| qtl4.1 | 4 | 1.96 | 15.119 | 1.3 | 1 | 5.9 | 0.3 | 0.8 | 0.3 |
| qtl4.2 | 4 | 5.41 | 35.609* | 2.6 | 0.1 | 6.8 | 10.4 | 0.8 | 0.4 |
| qtl4.3 | 4 | 11.16 | 65.482 | 2.6 | 0.4 | 6.2 | 2.4 | 0.3 | 0.9 |
| qtl4.4 | 4 | 49.49 | 78.122 | 0.3 | 0.7 | 2.1 | 6 | 0 | 0 |
| qtl4.5 | 4 | 49.86 | 79.403 | 1.3 | 1 | 0.7 | 0.5 | 5.3 | 0.1 |
| qtl4.6 | 4 | 51.88 | 84.269* | 1 | 0.6 | 5 | 0.9 | 6.3 | 1.3 |
| qtl4.7 | 4 | 58.07 | 107.048 | 0.4 | 1.2 | 5.6 | 0.5 | 0.2 | 0.1 |
| qtl4.8 | 4 | 61.65 | 126.443* | 1 | 0.1 | 1.7 | 0.1 | 9.1 | 9.6 |
| qtl4.9 | 4 | 63.79 | 139.512 | 0.4 | 0.2 | 0.6 | 6.3 | 0 | 2 |
| qtl4.10 | 4 | 66.08 | 155.624 | 2.4 | 7.7 | 1.2 | 0.5 | 0.1 | 0 |
| qtl5.1 | 5 | 2.01 | 15.124* | 6.4 | 0.2 | 8.2 | 0.5 | 1.4 | 1.5 |
| qtl5.2 | 5 | 4.3 | 24.776 | 0.9 | 0.7 | 0.6 | 2 | 0.1 | 5.4 |
| qtl5.3 | 5 | 7.3 | 45.175 | 0.7 | 0.7 | 0.1 | 0.6 | 12 | 0.9 |
| qtl5.4 | 5 | 52.58 | 71.611 | 0 | 1.3 | 0.4 | 0.1 | 5.8 | 1.2 |
| qtl5.5 | 5 | 64.15 | 124.193 | 0.2 | 3 | 0.4 | 1.4 | 1.6 | 8.2 |
| qtl5.6 | 5 | 67.3 | 130.294 | 6 | 1.2 | 4.4 | 1.8 | 1.1 | 0 |
| qtl5.7 | 5 | 70.9 | 137.263 | 0.2 | 5.9 | 1.5 | 0.3 | 0.4 | 0.9 |
| qtl6.1 | 6 | 1.01 | 8.287* | 8 | 1.1 | 7.8 | 1.3 | 0.4 | 0.7 |
| qtl6.2 | 6 | 1.87 | 22.817 | 0 | 0.5 | 1.1 | 1.1 | 8.3 | 0.9 |
| qtl6.3 | 6 | 3.83 | 35.124 | 0.5 | 6.5 | 1.7 | 0.1 | 0.4 | 0 |
| qtl6.4 | 6 | 47.47 | 70.959* | 3.3 | 5.3 | 6.2 | 1.7 | 0.1 | 2.7 |
| qtl6.5 | 6 | 51.76 | 90.609 | 0.3 | 7.5 | 0.5 | 0.1 | 0.6 | 0.2 |
| qtl6.6 | 6 | 53.69 | 111.316 | 0.2 | 5.1 | 0.7 | 0.2 | 0.2 | 0.1 |
| qtl6.7 | 6 | 54.19 | 115.77 | 1.6 | 0.5 | 1.4 | 1.7 | 0.4 | 11.3 |
| qtl6.8 | 6 | 54.69 | 122.137 | 0.1 | 0 | 0.1 | 8.4 | 0 | 0.1 |
| qtl6.9 | 6 | 58.18 | 149.111 | 0.4 | 1 | 0.2 | 1.4 | 0.4 | 9.6 |
| qtl6.10 | 6 | 60.77 | 162.382 | 0.4 | 0.5 | 0 | 0.2 | 7.6 | 1.1 |
| qtl7.1 | 7 | 1.1 | 5.133 | 3.6 | 1.1 | 1.3 | 1.9 | 5.6 | 1 |
| qtl7.2 | 7 | 9.24 | 70.318 | 7.4 | 1.1 | 1.5 | 1.7 | 0.2 | 2.2 |
| qtl7.3 | 7 | 60.6 | 117.74 | 0.4 | 1.1 | 0.1 | 8.2 | 0 | 0.6 |
| qtl7.4 | 7 | 61.43 | 123.307* | 6.9 | 1.2 | 1.3 | 1.1 | 18.2 | 5.7 |
| qtl8.1 | 8 | 0.81 | 8.54 | 0.1 | 0.6 | 0.5 | 0.6 | 2.7 | 6.3 |
| qtl8.2 | 8 | 2.12 | 19.183 | 1.2 | 0.6 | 2.8 | 0.4 | 8.9 | 0.6 |
| qtl8.3 | 8 | 5.48 | 58.691 | 0.3 | 0.6 | 1.8 | 7.2 | 0.9 | 1.7 |
| qtl8.4 | 8 | 19.27 | 67.65* | 4.5 | 5.6 | 5.9 | 2.4 | 5.1 | 1.1 |
| qtl8.5 | 8 | 56.75 | 135.757 | 0.1 | 1.1 | 0 | 0.3 | 10.5 | 2.5 |
| qtl8.6 | 8 | 61.25 | 146.531 | 0.6 | 0.4 | 0.6 | 1.6 | 0.3 | 6.4 |
| qtl9.1 | 9 | 0.53 | 7.058 | 0.2 | 0.5 | 0.2 | 1.6 | 7.4 | 1.2 |
| qtl9.2 | 9 | 1.99 | 26.157 | 3.7 | 0.5 | 5.3 | 2.4 | 0.8 | 1.8 |
| qtl9.3 | 9 | 2.4 | 32.616 | 5.4 | 1 | 1.8 | 0.4 | 6.9 | 0.4 |
| qtl9.4 | 9 | 4.58 | 52.724 | 7.2 | 0.1 | 1.1 | 0.7 | 0.1 | 0 |
| qtl9.5 | 9 | 6.39 | 61.212 | 0.3 | 0 | 0.9 | 1 | 9.5 | 0.2 |
| qtl9.6 | 9 | 19.46 | 70.336 | 0 | 0.1 | 0.1 | 0.5 | 0.1 | 5.2 |
| qtl9.7 | 9 | 53.53 | 106.98 | 7.1 | 0.1 | 2.6 | 3.6 | 0.6 | 2.5 |
| qtl9.8 | 9 | 56.52 | 114.89 | 6.3 | 0.5 | 1.5 | 0.5 | 0.8 | 1.6 |
| qtl10.1 | 10 | 0.41 | 7.332 | 6.7 | 0.1 | 0.3 | 1.8 | 0.4 | 0.9 |
| qtl10.2 | 10 | 1.84 | 26.837 | 0.6 | 7 | 0.7 | 0 | 0.3 | 0 |
| qtl10.3 | 10 | 2.56 | 29.394* | 1.1 | 0.9 | 7.8 | 1.8 | 11.8 | 13.7 |
| qtl10.4 | 10 | 4.31 | 33.318 | 0.2 | 0.7 | 1 | 5.5 | 0 | 0.1 |
| qtl10.5 | 10 | 6.14 | 36.912 | 1.1 | 5.1 | 0.7 | 0.3 | 0.6 | 0.2 |
| qtl10.6 | 10 | 9.34 | 52.009 | 0.5 | 0.1 | 1.6 | 5.5 | 1.7 | 0 |
| qtl10.7 | 10 | 16.06 | 57.905 | 1.3 | 1.1 | 5.1 | 0.7 | 0.1 | 0.4 |
| qtl10.8 | 10 | 50.05 | 63.653 | 1.5 | 0.1 | 0.6 | 0.7 | 1.9 | 9.9 |
| qtl10.9 | 10 | 50.83 | 65.967 | 7.3 | 0.3 | 0.9 | 0.4 | 0 | 0.4 |
| qtl10.10 | 10 | 52.9 | 72.962 | 0 | 0.1 | 0.5 | 1 | 0.6 | 5.2 |
| qtl10.11 | 10 | 54.03 | 76.78 | 0.3 | 0.3 | 0.1 | 0.5 | **6.7** | 0.2 |
| qtl10.12 | 10 | 55.48 | 81.237 | **6.4** | 0.8 | 0.6 | 0.1 | 0.3 | 0.6 |
| qtl10.13 | 10 | 60.19 | 109.735 | 0.6 | **9.2** | 2.5 | 0.2 | 0.6 | 0.9 |

Values represent $-\log_{10}(P)$ of the peak SNP within each QTL window, defined as $\pm 1$ cM around the peak SNP based on the genetic map. **Bold black** indicates values exceeding the genome-wide significance threshold ( $P < 1 \times 10^{-5}$ ; $-\log_{10} P > 5$ ). Cell colour intensity reflects the $-\log_{10}(P)$ value across all six analyses, allowing visual assessment of signal concordance across years and analysis types. Position (Mb) and Peak (cM) = physical and genetic map positions of the peak SNP, respectively. BL = baseline tillering analysis (isolated plants, no neighbour effect); R = neighbour-responsive tillering analysis (genotypes filtered for No < 1). Suffixes \_23 and \_24 denote 2023 and 2024 single-year analyses; \_com denotes the combined two-year analysis. Scale: 0 12. Scale values denote $-\log_{10}(P)$ magnitude (0-2: no signal; 2-4: background; 4-5: sub-threshold but elevated; 5-8: significant; 8+: highly significant). Sub-threshold values (4-5) are reported to show concordance of signal across analyses but were not counted as detected significant QTLs in that analysis.
\* Asterisk on Peak (cM) indicates the QTL window contains two distinct significant SNPs; only the position of the more significant SNP is shown here. Individual SNP positions, alleles, and $P$ -values for all significant SNPs are provided in Supplementary Table S3.

To evaluate the biological relevance of density-responsive tillering QTLs, we examined the enrichment of R:FR light signalling candidate genes within identified QTL regions. A set of 41 R:FR pathway candidate genes known to regulate neighbour detection and shade avoidance responses in plants were compiled from Arabidopsis and rice (Table S1). These candidate genes represent key components of neighbour detection mechanisms, including phytochrome signalling (*SbPhyA*, *SbPhyB*), phytochrome interacting factors (SbPIF5, SbPIF8), light signal transduction (*SbHY5*, *SbCOP1*, *SbBBX4*, *SbBBX11*, *SbBBX21*), shade avoidance (*SbSAV4*, *SbFHY1*, *SbSPA1*), and R:FR-regulated branching pathways (*SbMAX2*, *SbMAX4*, *SbBRC1*) (Ballaré *et al.,* 1987; Pierik & de Wit, 2014; Casal & Fankhauser, 2023).

Baseline tillering QTLs showed no enrichment for R:FR candidate genes, with zero genes falling within the peak SNP windows of the 52 QTLs regions. In contrast, the No<1 responsiveness analysis revealed significant enrichment of R:FR pathway genes. Five candidate genes co-localized with responsiveness QTLs: *SbPIF5* (Sobic.001G068301) within qtl1.3, *SbMAX4*_a (Sobic.003G293600) within qtl3.8, *SbBBX21*_a (Sobic.004G301000) within qtl4.9, *SbMAX4*_d (Sobic.007G170300) within qtl7.3, and *SbMAX2* (Sobic.010G043000) within qtl10.4, representing 10% of responsiveness QTLs (5/50) and 12.2% of candidate genes (5/41). No candidate genes co-localized with baseline tillering QTLs (0/41). Notably, two independent paralogues of MAX4 on different chromosomes (*SbMAX4*_a on chromosome 3 and *SbMAX4*_d on chromosome 7) each co-localized with separate responsiveness QTL, providing independent support for the involvement of R:FR-regulated branching pathways in density-responsive tillering. This differential enrichment was statistically significant (Fisher’s exact test, *P* = 0.025), indicating that responsiveness QTLs are specifically enriched for R:FR light signalling pathway genes.

## 4. Discussion

### 4.1. Methodology validation

This study introduces several methodological innovations that significantly advanced the genetic analysis of tillering plasticity in grasses. The mathematical model incorporating nearest (N) and farthest (F) neighbour distance effects (*Tiller* = *β*_0_ + *G* + *β*_0 1_*N* + *β*_0 2_*F* + *β*_0 3_(*N* x *F*) + *β*_0_ ₄*N*² + β₅*F*² + *e)* represents the first quantitative framework for capturing spatial neighbour detection mechanisms in GWAS studies (Hambäck *et al.,* 2014). Unlike traditional density treatments that rely on categorical comparisons, this approach quantifies how plants respond to neighbour at multiple spatial scales, with interaction and quadratic terms allowing for non-linear responses. Model fitness evaluation across both growing seasons demonstrated robust performance, with significant genotype-by-distance interactions (z-ratios > 2.0), confirming successful capture of genetic variation in neighbour responsiveness.

The systematic filtering strategy focusing on genotypes with sufficient baseline tillering capacity (No<1 filter) to detect response to density analysed 729 genotypes in 2023 and 545 genotypes in 2024. Using a 2-cM clustering approach (Tao *et al.,* 2020), the 65 significant SNPs clustered into 50 unique responsiveness QTLs, with only 10 instances of overlap with baseline tillering associations. Some overlap is expected as R:FR-responsive genes would also be active in isolated plants through self-shading effects within the plant canopy. The predominantly non-overlapping QTLs support successful identification of loci specifically involved in plastic neighbour-responsive mechanisms. This represents the first genome-wide association study in grasses to explicitly parameterise tillering responsiveness to plant spacing, establishing a framework that could be applied to future studies of phenotypic plasticity of tillering in crop species.

### 4.2. Genetic architecture of tillering and its plasticity

The absence of R:FR candidate gene enrichment in baseline tillering QTLs, despite these QTLs spanning approximately 7% of the sorghum genome, is a notable finding. This suggests that constitutive tillering capacity is predominantly governed primarily by developmental and hormonal pathways rather than light quality perception mechanisms. Although R:FR-responsive genes are likely expressed at some level in isolated plants through self-shading within the plant canopy, their contribution to genetic variation in baseline tillering appears insufficient for GWAS detection. The contrast between zero R:FR gene enrichment in baseline QTLs and significant enrichment (12.2%, 5/41 genes, Fisher’s exact test, *P* = 0.025) in responsiveness QTLs provides strong evidence that our filtering approach successfully separated two distinct genetic components of tillering: constitutive developmental control and plastic neighbour-responsive control.

This study reveals that tillering capacity and tillering plasticity in sorghum are controlled by shared and interconnected genetic architectures. Baseline tillering capacity, representing the inherent ability to produce tillers without competitive pressure, involves at least 52 detectable QTLs distributed across all 10 chromosomes, demonstrating the highly polygenic nature of constitutive tillering mechanisms. Based on previous studies in cereals, constitutive tillering is known to involve developmental processes including axillary bud activation, phytohormone signalling pathways, and sugar sensing mechanisms (Li *et al.,* 2003; Domagalska & Leyser, 2011; Kebrom *et al.,* 2013; Mason *et al.,* 2014). Although we did not test enrichment for these specific pathway genes among our baseline QTLs, the finding that baseline QTLs contain none of the 41 R:FR candidate genes suggests that constitutive tillering is controlled by different genetic mechanisms than neighbour-responsive tillering. The 20% overlap between baseline and responsiveness QTLs suggests some shared genetic control between constitutive and plastic tillering. Notably, none of the five R:FR candidate genes identified in responsiveness QTLs were located within these overlapping regions, indicating that the shared QTLs may involve regulatory mechanisms other than the R:FR light signalling pathways detected in this study. Self-shading, where a plant’s own canopy alters the R:FR ratio experienced by lower buds, could contribute to this overlap, though further characterisation of genes within overlapping QTL regions would be needed to test this hypothesis.

However, tillering plasticity represents a specialized subset of this broader genetic architecture, involving 50 QTL of which 10 overlap with baseline tillering QTLs. The identification of 10 QTLs that overlap between baseline tillering and neighbour responsiveness confirms the shared regulatory mechanisms controlling both basic and plastic aspects of tillering behaviour. The contrasting genetic architectures between baseline tillering (broadly distributed, polygenic) and plasticity (focused, specialized) indicate that evolutionary selection has shaped distinct molecular mechanisms for constitutive growth versus competitive responses (Ballaré *et al.,* 1994; Weiner, 1990).

The limited overlap in QTL detection between years can be attributed to two interconnected factors: the complexity of the trait and reduced statistical power from our filtering methodology and genotype × environment (GxE) interactions. While the density-responsiveness filtering strategy was supported through enrichment of R:FR pathway genes, it necessarily reduced sample sizes from 895 individuals to smaller subsets of genotypes where responsiveness could be measured. Importantly, these subsets differed between years due to environmental variation, creating distinct environments for analysis. The combination of reduced statistical power caused by the complex genetic architecture of tillering and the inherently environment-sensitive nature of tillering plasticity responses resulted in year-specific QTLs detection under the contrasting environmental conditions of 2023 and 2024. Rather than representing a methodological limitation, these year-specific QTLs are interpreted to reflect genuine genetic regulation of neighbour-responsive tillering. This finding is consistent with the adaptive nature of phenotypic plasticity and indicates that our filtering approach captured genetic mechanisms associated with plastic tillering responses successfully.

### 4.3. QTL co-localization with red:far-red light signalling pathways

The enrichment of red and far red light signalling and phytochrome pathway genes among density-responsive QTLs provide strong mechanistic evidence for neighbour detection through R:FR light ratio changes (Smith, 1995; Ballaré *et al.,* 1997). This co-localization indicates that sorghum, like other grasses and dicots, utilises conserved shade avoidance mechanisms for regulating tillering responses to neighbour competition. The molecular basis involves detection of altered spectral quality when neighbouring plants reflect far-red light while absorbing red light, creating reliable signals of competitive intensity (Holmes & Smith, 1977; Casal, 2012). Several major responsiveness QTL regions contain candidate genes involved in phytochrome signal transduction and downstream regulatory cascades that modify developmental programs in response to competitive intensity (Franklin, 2008). Beyond this enrichment, the unique responsiveness QTLs analysis could reveal evidence of further regulatory gene network involvement in plasticity. These QTLs provide targets for further experimental investigations. The five co-localized R:FR pathway genes represent two interconnected branches of the phyB signalling network. *SbPIF5* directly interacts with phyB and accumulates under low R:FR to promote shade avoidance responses including reduced branching (Shen *et al.,* 2007; Wu *et al*., 2019), while *SbBBX21* upregulates HY5 expression downstream of phytochrome-mediated inactivation of the COP1/SPA complex (Xu *et al.,* 2018). The remaining three co-localized genes, *SbMAX2*, *SbMAX4_a*, and *SbMAX4_d*, are components of the strigolactone pathway that is directly regulated by R:FR light. In sorghum, *SbMAX2* expression in axillary buds is elevated in *phyB-1* mutants and under far-red light supplementation (Kebrom *et al*., 2010), and phyB-dependent regulation of bud outgrowth requires functional *MAX4* (Finlayson *et al.,* 2010). Recent evidence demonstrates that FHY3 and FAR1, two phyA signalling components, directly integrate light signalling with the MAX2/MAX4 pathway to regulate branching through BRC1 (Xie *et al.,* 2020). The exclusive co-localization of both direct light perception components and R:FR-regulated strigolactone genes with responsiveness QTLs, and their complete absence from baseline QTLs, demonstrates that the No<1 filtering approach effectively captured genotypes with authentic tillering plasticity responses. This result provides support that the density-responsive QTLs detected represent genetic loci involved in known mechanisms controlling tillering plasticity, supporting both our mathematical modelling approach and filtering methodology for quantifying neighbour responsiveness in sorghum.

Cross-species comparative analysis reveals that the phytochrome pathway genes identified in sorghum likely have functional orthologs in maize and rice, where similar tillering plasticity mechanisms could be targeted to improve yield stability in high-density planting systems (Whipple *et al.,* 2011; Xu *et al.,* 2012). The conservation of light-mediated neighbour detection pathways across species is well established (Chen *et al.,* 2013; Hirose *et al.,* 2012), and our identification of conserved phytochrome pathway genes within sorghum responsiveness QTL is consistent with this interpretation. The identification of both conserved and novel QTLs suggest that tillering plasticity may involve a combination of shared regulatory mechanisms and lineage-specific genetic effects, consistent with previous studies of plant competition (Ballaré *et al.,* 1994; Weiner, 1990), and connecting fundamental research on plant competition with practical application of crop improvement.

### 4.4. Implications for crop improvement and future applications

Tillering plasticity is a crucial adaptive trait that enables sorghum plants to optimize their architecture in response to competitive environments, representing a fundamental mechanism for balancing resource allocation between main culm development and tiller production under varying densities. While this trait provides clear advantages in enhancing survival and competitiveness in natural ecosystems, its utility in agricultural systems requires more careful consideration. In modern sorghum farming systems establishment and plant spacing are highly consistent and plant responses to spacing may have negative consequences in some circumstances. For example, in water-limited environments, farmers can utilize information about soil moisture at planting, climate forecasting combined with precision planting to optimize canopy size and water use for the expected season (Durrington *et al.,* 2025). Under such circumstances, farmers would benefit from predictable plant architectures tailored to the expected specific environment at key development stages rather than relying on natural plasticity responses. For example, farmers may target a smaller canopy size by planting at low density within rows or wider row spacing when they expect water-limited conditions to avoid the severe yield reductions associated with terminal drought stress (Hammer *et al.,* 2023). Under these circumstances, genotypes that do not respond to density are preferred as they enable changes in density and row spacing to directly change canopy size and water use, whereas plastic varieties will respond to low spacing by increasing tillering, canopy size and water use. The genetic diversity uncovered in our study presents valuable opportunities for crop improvement through marker-assisted selection or genomic prediction and selection to produce varieties more suited to modern management. Given the polygenic nature of tillering responsiveness revealed in this study, genomic prediction incorporating these QTLs as features would likely be a more effective breeding strategy than targeting individual loci through marker-assisted selection. The identification of specific pathway genes such as *PIF5* and *BBX21* within responsiveness QTLs provides valuable biological insight that could inform genomic prediction models and prioritise candidate loci for functional validation, ultimately contributing to the development of cultivars with either controlled responsiveness or stable architectures suited to specific environmental conditions. This approach would allow separating the natural plasticity mechanism from yield potential, enabling the development of sorghum varieties better adapted to modern agricultural systems where environmental predictability often outweighs the advantages of natural plasticity mechanisms.

The broader implications of this work extend beyond sorghum improvement. The conserved nature of light-mediated neighbour detection suggests that similar QTL mapping approaches could be applied to other grass crops including wheat and rice, offering opportunities for cross-species breeding strategies. Beyond immediate applications for sorghum breeding, these findings contribute to fundamental understanding of how plants have evolved mechanisms for detecting and responding to neighbour competition. As agriculture increasingly requires crops with enhanced phenotypic plasticity to maintain stable production under variable conditions, this study provides both the conceptual framework and molecular insights necessary for achieving this potential in sorghum and related grass crops.

## Acknowledgements

This research was supported by the Australian Research Council Centre of Excellence for Plant Success in Nature and Agriculture (grant no.CE200100015). AR was supported by a University of Queensland Research Training Scholarship. We thank the field staff at Hermitage Research Facility for technical support with field trials.

## Author contributions

DJ and EM conceived and designed the study. AR conducted the field trials, performed phenotyping, statistical analysis and GWAS, and drafted the manuscript. SP assisted analysis and reviewing the manuscript. CH provided statistical guidance and contributed to the spatial modelling approach. SS contributed to data collection and reviewing manuscript. YT contributed to analysis and reviewing manuscript. MC and GH contributed to project conceptualisation and interpretation of results in the context of crop modelling. EM and DJ supervised the project and provided manuscript editing. All authors reviewed and approved the final manuscript.

## Data availability

Data available in article Supporting Information (Figure S1; Tables S1-S3).

